# Phylogenomic Testing of Root Hypotheses

**DOI:** 10.1101/758581

**Authors:** Fernando D. K. Tria, Giddy Landan, Devani Romero Picazo, Tal Dagan

## Abstract

The determination of the last common ancestor (LCA) of a group of species plays a vital role in evolutionary theory. Traditionally, an LCA is inferred by the rooting of a fully resolved species tree. From a theoretical perspective, however, inference of the LCA amounts to the reconstruction of just one branch - the root branch - of the true species tree, and should therefore be a much easier task than the full resolution of the species tree. Discarding the reliance on a hypothesised species tree and its rooting leads us to re-evaluate what phylogenetic signal is directly relevant to LCA inference, and to recast the task as that of sampling the total evidence from all gene families at the genomic scope. Here we reformulate LCA and root inference in the framework of statistical hypothesis testing and outline an analytical procedure to formally test competing *a-priori* LCA hypotheses and to infer confidence sets for the earliest speciation events in the history of a group of species. Applying our methods to two demonstrative datasets we show that our inference of the opisthokonta LCA is well in agreement with the common knowledge. Inference of the proteobacteria LCA shows that it is most closely related to modern Epsilonproteobacteria, raising the possibility that it may have been characterized by a chemolithoautotrophic and anaerobic life-style. Our inference is based on data comprising between 43% (opisthokonta) and 86% (proteobacteria) of all gene families. Approaching LCA inference within a statistical framework renders the phylogenomic inference powerful and robust.

## Introduction

Inference of the last common ancestor (LCA) for a group of taxa is central for the study of evolution of genes, genomes and organisms. Prior to the assignment of an LCA, represented by the root node, phylogenetic trees are devoid of a time direction and the temporal order of divergences is undetermined. Discoveries and insights from the reconstruction of LCAs span a wide range of taxonomic groups and time scales. For example, studies of the last universal common ancestor (LUCA) of all living organisms inferred that it was an anaerobic prokaryote whose energy metabolism was characterized by CO_2_-fixing, H_2_-dependent with a Wood– Ljungdahl pathway, N_2_-fixing and thermophilic (Weiss et al. 2016). Nonetheless, due to the inherent difficulty of ancient LCA inference, it is frequently the centre of evolutionary controversies, such as for instance the debate concerning the two versus three domains of life (Williams et al. 2020), the cyanobacterial root position (Hammerschmidt et al. 2021), the LCA of vertebrates (Okamoto et al. 2017), or the LCA of hominids (Lovejoy et al. 2009).

The identity of the LCA is traditionally inferred from a species tree that is reconstructed unrooted and is then rooted at the final step. The root may be inferred by several methods, including: outgroup rooting (Kluge and Farris 1969), midpoint rooting (Farris 1972), minimal ancestor deviation (Tria et al. 2017), Minimal variance rooting (Mai et al. 2017), relaxed clock models (Lepage et al. 2007), tree reconciliation (Szöllősi et al. 2012), as well as non-reversible substitution models for nucleotide data (Huelsenbeck et al. 2002; Williams et al. 2015; Cherlin et al. 2018; Bettisworth and Stamatakis 2021) or amino acids data (Naser-Khdour et al. 2022). A correct LCA inference in methods that rely on the availability of the species tree, is thus dependent on the accuracy of the species tree topology. One approach for the reconstruction of a species tree, is to use a single gene as a proxy for the species tree topology, e.g., 16S ribosomal RNA subunit for prokaryotes (Fox et al. 1980) or the Cytochrome C for eukaryotes (Fitch and Margoliash 1967). This approach is, however, limited in its utility due to possible differences between the gene evolutionary history and the species phylogeny. Similarly, methods relying on ancient paralogs for root inferences (e.g., Gogarten et al. 1989; Iwabe et al. 1989) rely on a small number genes and assume an identity of the gene and species evolutionary history.

Phylogenomics offer an alternative to the single-gene approach by aiming to utilize the whole genome rather than a single gene for the phylogenetic reconstruction (Eisen 2003). In the most basic approach, the species tree is reconstructed from the genes that are shared among all the species under study, termed here *complete gene families.* Approaches for the reconstruction of a species tree from multiple single-copy genes include tree reconstruction from concatenated alignments (e.g., Ciccarelli et al. 2006; Hug et al. 2016; Parks et al. 2018) as well as calculation of consensus trees (e.g., Dagan et al. 2013). However, these approaches are often restricted in their data sample as they exclude *partial gene families* not present in all members of the species set (e.g., due to differential loss) and *multi-copy gene families* present in multiple copies in one or more species (e.g., as a result of gene duplications or gene acquisition). Thus, the drawback in alignment concatenation or consensus tree approaches is that the inference becomes limited to gene sets that do not represent the entirety of genomes. This issue tends to become more acute the more diverse the species set is, or when it includes species with reduced genomes. In extreme cases no single-copy and complete gene family exists (Medini et al. 2005). Super-tree approaches offer an alternative as they enable to include also partial gene families (Pisani et al. 2007; Whidden et al. 2014; Williams et al. 2017); however, those approaches also exclude multi-copy gene families (partial and complete). These requirements produce the phenomenon of “trees of 1%” of the genes (Dagan and Martin 2006; and see our Table 1). Thus, while the major aim of phylogenomics approaches is to improve the accuracy of phylogenetic inference by increasing the sample size, methodologies based on single-copy genes suffer from several inference problems, with some elements that are common to all of them. The first is the limited sample sizes due to the number of single copy genes. In super-tree approaches based on concatenation and consensus there is no room for the inclusion of multi-copy gene families. Furthermore, reduction of families including paralogs into orthologs-only subsets, e.g., using tree reconciliation (reviewed in (Szöllősi et al. 2015; Smith and Hahn 2021)), requires an a-priori assumed species tree topology. Finally, since the aforementioned approaches yield unrooted species trees, the inference of the root is performed as the last step in the analysis, hence the sample size of trees used for the LCA inference is essentially one (i.e., a single tree).

The inference of the LCA from a single species tree can be robust and accurate only if the underlying species tree is reliable. Unfortunately, this is rarely the case, as can be frequently seen in the plurality of gene tree topologies and their disagreement with species trees, especially for prokaryotes due to frequent gene transfers (e.g., Doolittle and Bapteste 2007; Linz et al. 2007; and see our Fig. 5a). Indeed, recent implementations of tree reconciliation approaches aim to accommodate the presence of heterogeneous topologies due to gene duplication or gain by inferring the effect of such events on the tree topology (e.g., (Coleman et al. 2021; Morel et al. 2022)). However, such applications required an *a-priori* assumption of the relative frequency of gene duplication and gene transfer, that is bound to have a significant effect on the resulting tree topology (including the root position) (Bremer et al. 2022). We propose that the identification of an LCA for a group of species does not require the reconstruction of a fully resolved species tree. Instead, the LCA can be defined as the first speciation event, i.e., tree node, for the group of species. In this formulation, the topological resolution of the entire species tree is immaterial and the only phylogenetic conclusion needed is the partitioning of the species into two disjoint monophyletic groups or – the *species root partition.*

Here we present a novel approach for the LCA inference without reconstructing a species tree. Our approach considers the total evidence from unrooted gene trees for all protein families from a set of taxa, including partial families as well as multiple-copy gene families. The approach utilizes the measure of ancestor deviation (AD) that is the basis for the minimal ancestor deviation (MAD) rooting method (Tria et al. 2017). The MAD rooting method assumes a clock-like evolutionary rate of protein families, which has been shown to be a reasonable assumption, at least for prokaryotic protein families where ca. 70% families do not deviate significantly from clock-like evolutionary rate (Novichkov et al. 2004; Dagan et al. 2010). The AD measure is calculated for each branch in a given phylogenetic tree as the mean relative deviation from the molecular clock expectation, when the root is positioned on that branch. The branch that minimizes the relative deviations from the molecular clock assumption (i.e., is assigned the minimal AD) is the best candidate to contain the root node. The AD calculation can be performed for any given tree topology, regardless of the tree reconstruction approach. The comparison of branch ADs calculated for gene trees of all protein families in a group of species enables us to formulate and test hypotheses regarding the LCA of the species tree.

## Results

For the presentation of the statistical framework for testing root hypotheses, we used illustrative rooting problems for two species sets: opisthokonta and proteobacteria (Table 1). The root partition is well established for the opisthokonta species sets, and it serves here as a positive control. The root of the proteobacteria species set is still debated, hence this dataset serves to demonstrate the power of the proposed approach. The opisthokonta dataset comprises 14 metazoa and 17 fungi species, and the known root is a partition separating fungi from metazoa species (Stechmann and Cavalier-Smith 2002; Katz 2012). The proteobacteria dataset includes species from five taxonomic classes in that phylum (Ciccarelli et al. 2006; Pisani et al. 2007; Lang et al. 2013; but see Waite et al. 2017). The Proteobacteria dataset poses a harder root inference challenge than opisthokonta as previous results suggest that three different branches are comparably likely to harbour the root node, a situation best described as a *root neighbourhood* (Tria et al. 2017).

**Table 1.** Illustrative datasets, their consensus rooting using minimal ancestor deviation (MAD), and classification of gene families. For the complete list of species, see Supplementary Table 1.

|  | Opisthokonta | Proteobacteria |
| --- | --- | --- |
| Number of species | 31 | 72 |
| **Consensus MAD Root in CSC gene trees** | 77.70%<br>(132 trees)<br>{Fungi, Metazoa} | 33.17%<br>(16,583 trees)<br>{ε-proteobacteria, other proteobacteria} |
| **Number of gene families** | 13,036 | 9,686 |
| **CSC:** Complete single-copy gene families, present as single-copy in all members of a species set. | 170<br>(1.30%) | 50<br>(0.52%) |
| **CMC:** Complete multi-copy gene families, present in all species, but having multiple copies in at least one species. | 612<br>(4.69%) | 70<br>(0.72%) |
| **PSC:** Partial single-copy gene families, absent from some species and present as single-copy in the others. | 7,773<br>(59.63%) | 5,586<br>(57.67%) |
| **PMC:** Partial multi-copy gene families, absent from some species and having multiple copies in at least one other species. | 4,481<br>(34.37%) | 3,980<br>(41.09%) |

### Phylogenomic Rooting as Hypothesis Testing

Our LCA inference approach differs from existing ones in several aspects: 1) No species tree is reconstructed or assumed. 2) Phylogenetic information is extracted from gene trees reconstructed from partial and multi-copy gene families in addition to CSC gene families. 3) The analysis uses unrooted gene trees and no rooting operations are performed, of either gene trees or species trees. 4) Any LCA hypothesis can be tested, including species partitions that do not occur in any of the trees. LCA inference deals with abstractions of similar but distinct types of phylogenetic roots: species tree roots and gene tree roots. We reserve the terms *root branch* or *root split* to refer to gene tree roots, while *species root partition* refers to species trees.

Before describing our approach, we first demonstrate the limitations of a simpler phylogenomic rooting procedure that uses only CSC gene families and infers the root by a consensus derived from the rooted trees of the CSC genes. In this procedure, only the root branch in each gene tree is considered for the root inference. We then show how to consider all branches from each unrooted gene tree, not only the root branch of the rooted trees. The incorporation of all branches, not considered by a simple consensus of rooted trees, leads to a statistical test to decide between two competing root hypotheses.

Next, we show how information from partial and multi-copy gene families can be used within the same statistical framework, greatly increasing the sample size and inference power. We then extend the pairwise formulation and consider multiple competing root partitions by testing all partition pairs, one pair at a time.

Finally, we modify the pairwise test to a comparison of one root partition against all alternatives simultaneously (a one-to-many test), and present a sequential elimination process that infers a minimal root neighbourhood, i.e., a confidence set of LCA partitions.

### Phylogenomic consensus rooting

The consensus approach infers the root partition of a species set from a sample of rooted CSC gene trees. Root branches are collected from all trees and the OTU split induced by the most frequent root branch in the sample is the inferred species root partition for the species set. In species sets with a strong root signal, this majority-rule approach is sufficient to determine a clear root partition for the species set. This circumstance is observed in the opisthokonta illustrative dataset. Using MAD (Tria et al. 2017) to root the individual gene trees, the consensus species root partition was inferred as the root branch found in more than 70% of the CSC gene trees (see Table 1). In the proteobacteria, in contrast, the most frequent root branch was inferred in just 33% of the CSC gene trees, while two competing root branches are observed in almost 15% of the gene trees each. The performance of the consensus approach is thus hindered by three factors. First, majority-rule voting considers just one branch from each gene tree, ignoring a large measure of the phylogenetic signal present in the gene trees. In addition, the quality of the root inference varies among the gene trees and is quantifiable, but this information is not utilized by the consensus approach. Lastly, simple voting cannot be satisfactorily tested for statistical significance.

### The root support test for two alternative root partitions

The first step in our approach is a formulation of a test to select between two competing species root partitions (see Fig. 1 for a road-map of the procedure). To that end, we do not infer a single root for each gene tree, but consider every branch of a gene tree as a possible root branch, and assign it a score that quantify the relative quality of different root positions. In the current study we use the Ancestor Deviation (AD) measure to assess the relative strength of alternative roots of the same gene tree. The AD statistic quantifies the amount of lineage rate heterogeneity that is induced by postulating a branch as harbouring the root of the tree (a detailed definition of AD is presented in the Appendix). We have previously shown that AD measures provide robust evidence for the inference of the root of a single gene tree (Tria et al. 2017).

**Figure 1.**
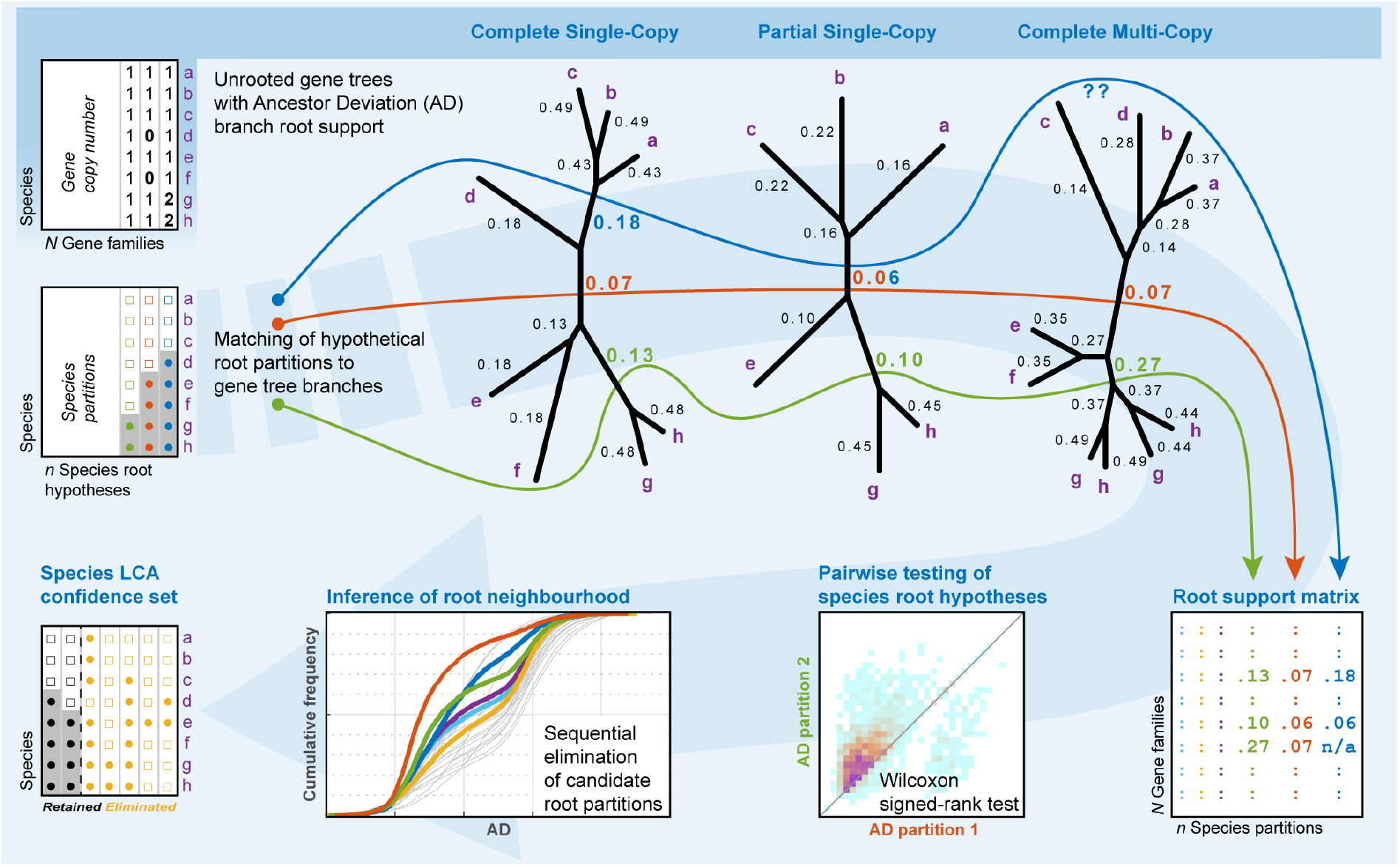
Outline of the analytical procedure. Stages are depicted clockwise from top-left. The input for the analysis is the gene trees of all protein families for a group of species, including the information of AD per branch as calculated by MAD. Protein families are classified into complete and partial, single- or multi-copy families according to the gene copy number per species. Branch ADs in the gene trees supply evidence for hypothetical root partitions in the species tree; these are collected in the root support matrix. The information in the root support matrix is used to identify candidates for the species root partition (including the consensus root partition, if exists). The comparison of root candidates is done by comparing the distribution of their ADs in all gene trees in a pairwise test. If several root partitions are similarly supported by ADs, these can be analysed in the context of a root neighbourhood, where weakly supported partitions are sequentially eliminated from the root partitions set. The remaining root partitions comprise the species LCA confidence set.

Collecting AD values from a set of gene trees, we obtain a paired sample of support values. In Fig. 2 we present the joint distribution of AD values for the two most likely root partitions in the eukaryotic dataset. A null hypothesis of equal support can be tested by the Wilcoxon signed-rank test, and rejection of the null hypothesis indicates that the root partition with smaller AD values is significantly better supported than the competitor (Fig. 2b).

**Figure 2.**
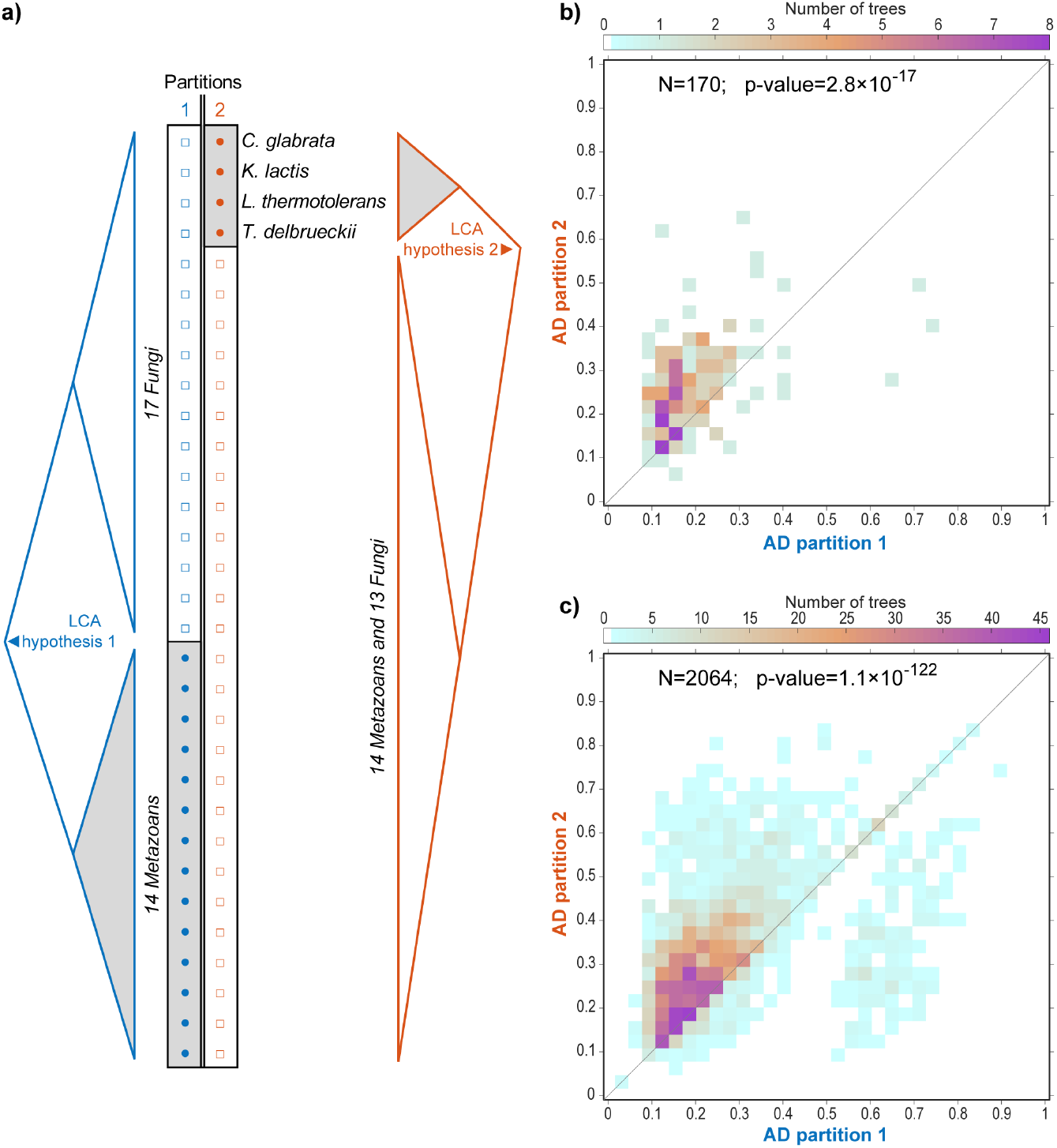
Pairwise testing of competing root hypotheses in the opisthokonta dataset. **a)** The two most frequent root branches among the CSC gene trees (Supplementary Table 1a). **b)** CSC gene families; **c)** All gene families. Colormaps are the joint distribution of paired AD values. Smaller ADs indicate better support, whereby candidate 1 out-compete candidate 2 above the diagonal and candidate 2 wins below the diagonal. p-values are for the two-sided Wilcoxon signed-rank test used to compare paired branch AD values. Note the gain in power concomitant to larger sample size.

As in all statistical inferences, the power of the test ultimately depends on the sample size. Considering only CSC gene families often limits rooting analyses to a small minority of the available sequence data (e.g., Table 1). Paired AD support values, however, can be extracted also from partial and multi-copy gene families, resulting in much larger sample size and statistical power (Fig. 2c).

### Rooting support from partial and multi-copy gene trees

To deal with non-CSC gene trees we must decouple the notion of *OTU split* (i.e, tree branch) from the notion of *species partition.* In CSC gene trees the correspondence between tree branches and root partitions is direct and one-to-one (Fig. 3a). In trees of partial gene families, a single OTU split may correspond to several species partitions. In multiple-copy gene families, some tree branches do not correspond to any possible species partition.

**Figure 3.**
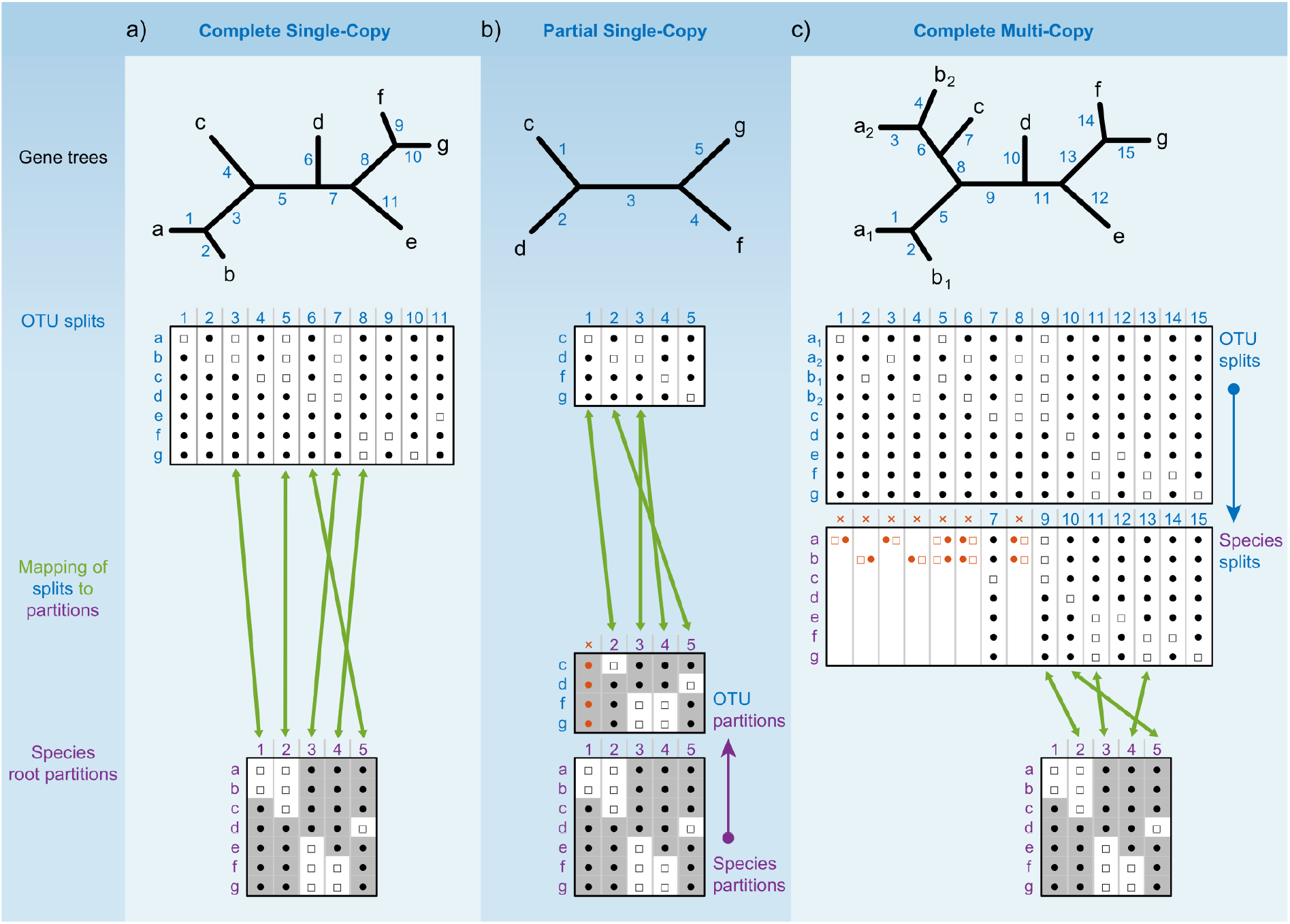
Correspondence of OTU splits and tested root partitions. **a)** In CSC gene trees; **b)** In PSC gene trees; **c)** In CMC gene trees. PMC gene trees entail both the **b)** and **c)** operations. OTU splits refer to branches in gene trees and are represented as black circles and white squares. Species partitions refer to possible branches in the hypothetical species tree, with unknown topology, and are represented as grey shades. In CSC gene trees all branches (including internal and external) can be mapped to species partitions in a one-to-one manner (green arrows in **a**; note that only several splits are illustrated). For mapping branches from PSC gene trees (**b**) to species partitions we remove from the species partitions the species missing in the gene tree and term the reduced version of the species partitions as OTU partitions. In CMC gene trees (**c**) only branches that form *species splits* can be mapped onto species partitions. A *species splits* in a CMC gene tree is a branch for which all gene copies of any one species are present on the same side of the split.

In order to find the branches in a *partial* gene tree that correspond to the tested root partitions, we *reduce* root partitions from species to OTUs by removing the species that are missing in the gene tree (Semple and Steel 2001). The root partitions are then assigned AD support by matching their reduced OTU version to the OTU splits of the gene tree (Fig 3b).

In *multi-copy* gene trees one or more species are represented multiple times as an OTU (Swenson and El-Mabrouk 2012). Each branch of a multi-copy gene tree splits the OTUs into two groups, and the two groups may be mutually exclusive or overlapping in terms of species. We refer to mutually exclusive splits in multi-copy gene trees as *species splits* which can be mapped to specific root partitions. Overlapping splits, on the other hand, cannot correspond to any root partition (Figure 3c). Mapping of tree splits from partial multi-copy gene trees entail both operations: identification of species splits and reduction of root partitions. We note that the ability to gain information on root partitions from multi-copy gene trees depends on the quality of gene family clustering with regards to the accuracy of orthology assessment. The presence of paralogs (especially ancient paralogs) in the set of multi-copy gene trees will lead to a low proportion of splits that can be mapped to root partitions.

Candidate root partitions, or their reduced versions, may be absent from some gene trees, and will be missing support values from these trees. We distinguish between two such cases: informative and uninformative missing values. A gene family is uninformative relative to a species root partition when its species composition includes species from only one side of the species partition. In such cases, the candidate root partition cannot be observable in any reconstructed gene tree. We label the gene trees of such families as uninformative relative to the candidate root partition, and exclude them from tests involving that partition. In contrast, when a gene family includes species from both sides of a candidate species root partition but the gene tree lacks a corresponding branch, we label the gene tree as informative relative to the partition. This constitutes evidence against the candidate partition, and should not be ignored in the ensuing tests. In such cases we replace the missing support values by a pseudo-count consisting of the maximal (i.e., worst) AD value in the gene tree. This assignment of a default worst-case support value also serves to enable the pairwise testing of incompatible root partitions, where no gene tree can include both partitions (Semple and Steel 2001).

Complete gene families are always informative relative to any candidate root partitions. Partial gene families, however, may be uninformative for some root candidates. When testing two candidate root hypotheses against each other, the exclusion of uninformative partial gene trees thus leads to a reduction of sample size from the full complement of gene families. Furthermore, one branch of a partial gene tree may be identical to the reduced versions of two or more species root partitions, whereby the tree is informative relative to the several candidates yet their support values are tied.

### Root inference and root neighborhoods

The pairwise test is useful when the two competing root hypotheses are given *a-priori,* as often happens in specific evolutionary controversies. More generally, however, one wishes to infer the species LCA, or root partition, with no prior hypotheses. In principle, the pairwise test may be carried out over all pairs of possible root partitions, while controlling for multiple testing. Such an exhaustive approach is practically limited to very small rooting problems, as the number of possible partitions grows exponentially with the number of species. A possible simplification is to restrict the analysis to test only pairs of root partitions from a pool of likely candidates. We propose that a reasonable pool of candidate root partitions can be constructed by collecting the set of root splits that are inferred as the root in any of the CSC gene trees.

When one species root partition is significantly better supported than any of the other candidates, the root is fully determined. Such is the result for the opisthokonta dataset, for which the known root partition is the best candidate among all pairwise comparisons (Supplementary Table 2a). In more difficult situations the interpretation of all pairwise *p*-values is not straightforward due to the absence of a unanimous best candidate root partition. This situation is exemplified with the CSC subset of the proteobacteria dataset where no candidate has better support than all the alternative candidates (Figure 4 and Supplementary Table 2b). The absence of a clear best candidate suggests the existence of a root neighbourhood in the species set. Thus, a rigorous procedure for the inference of a confidence set for LCA is required.

**Figure 4.**
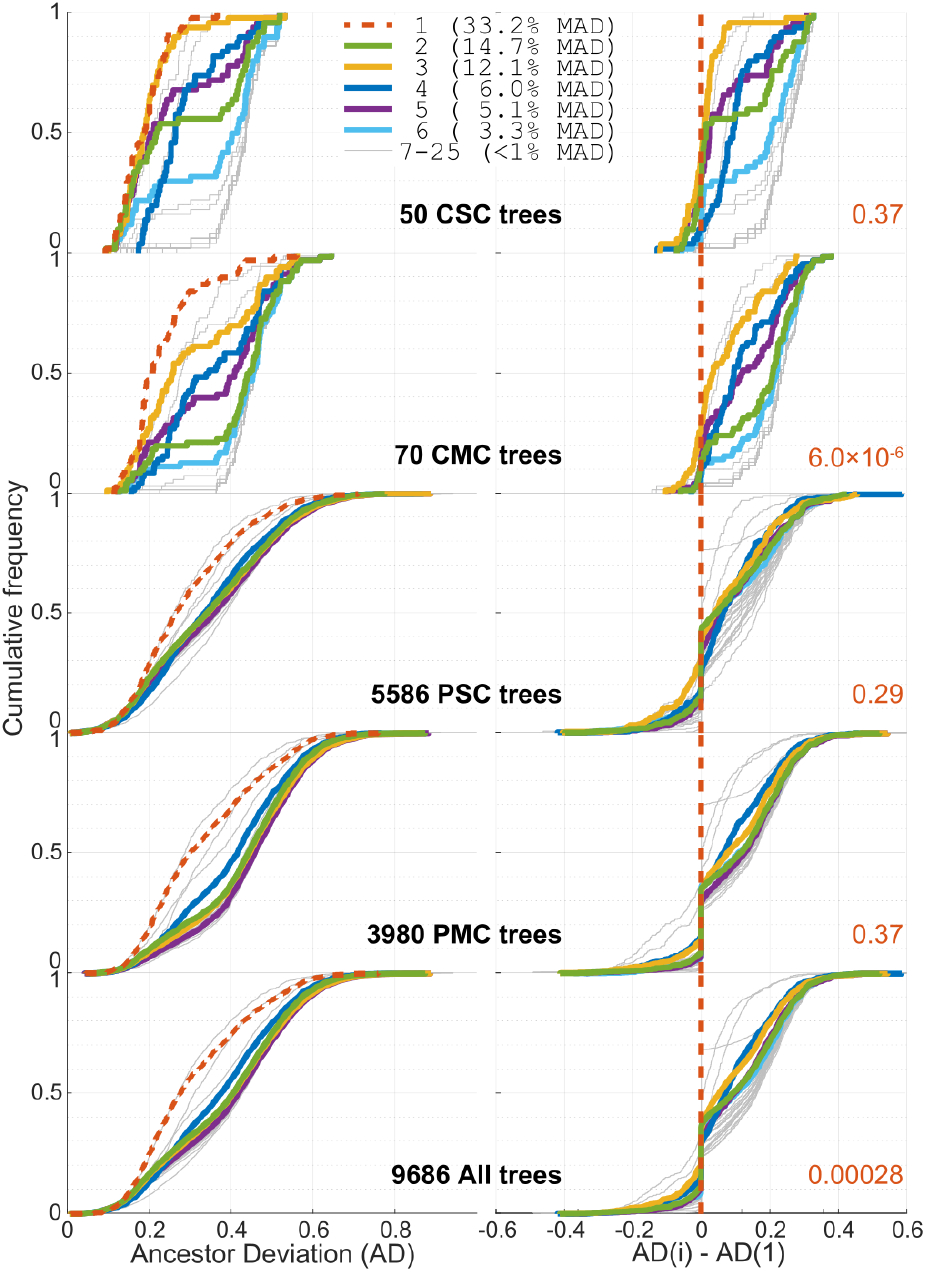
Cumulative distribution plots of AD in the proteobacteria dataset. Vertically are stacked the subsets by gene family type. On the left are unpaired AD values for the 25 candidate root partitions. On the right are the paired differences to candidate 1, whereby positive differences indicate better support for candidate 1 and negative values better support for candidate i. (See Supplementary Table 1b for candidate partition definitions). In red are p-values of the least significant among the contrasts to candidate 1, FDR adjusted for all 300 pairwise comparisons (Supplementary Table 2b).

### One-to-many root support test

To assess the support for root partitions in the full context of all other candidate root partitions, we modify the pairwise test to a test contrasting one root partition to a set of many alternatives. The *One-to-Many test* consists of comparing the distribution of root support values for one focal partition to the extreme support values among all the other candidates, and is inherently asymmetric. A ‘Better than Best’ version takes the minimal (i.e., best) value among the AD values of the alternatives, while the ‘Worse than Worst’ version considers the maximal (i.e., worst) among the alternatives’ ADs. As expected, the ‘better than best’ variant is always less powerful than any of the pairwise tests, and will not be considered further. The ‘worse than worst’ variant, on the other hand, can be used to trim down a set of candidates while being more conservative than the pairwise tests. In the one-to-many test, each gene tree provides one AD value for the focal partition and one AD value for the worst among the alternative root partitions. The worst AD values are assigned while considering only partitions with non-missing values (see above). Note that the worst alternative root partition may vary across gene trees. To maximize the sample size, i.e., number of trees used for the statistical tests, gene trees including informative missing values for all root candidate partitions are included in the analysis with the largest (i.e., worst) AD found in the entire tree. We test for differences in the magnitude of paired AD values using the one-sided Wilcoxon-signed rank test, with the null hypothesis that the focal ADs are equal or smaller than the maximal ADs for the complementary set, and the alternative hypothesis that the focal ADs are *larger still* than the maximum. A rejection of the null hypothesis is interpreted to mean that the focal root partition is significantly worse supported than the complementary set of candidates taken as a whole.

### Inference of a minimal root neighbourhood

To infer a root neighbourhood, i.e., a confidence set of LCA hypotheses, we start with a reasonably constructed large set of *n* candidate partitions, and reduce it by a stepwise elimination procedure. At each step, we employ the one-to-many test to contrast each of the remaining candidates to its complementary set. We control for false discovery rate (FDR; Benjamini and Hochberg 1995) due to multiple testing, and if at least one test is significant at the specified FDR level, the focal partition with the smallest p-value (i.e., largest z-statistic) is removed from the set of candidates. The iterative process is stopped when none of the retained candidates is significantly less supported than the worst support for the other members of the set, or when the set is reduced to a single root partition. To be conservative, we use a cumulative FDR procedure where at the first step we control for *n* tests, in the next round for *2n-1* tests, and, when not stopped earlier, for *n*(n-1)/2-1* at the last iteration.

We demonstrate the sequential elimination procedure for the proteobacteria dataset in figure 5. The splits network reconstructed for the proteobacteria dataset exemplifies the plurality of incongruent splits in the CSC gene trees and hence the dangers in assuming a single species tree. In this dataset the initial candidate set consisted of the 25 different root partitions found in the 50 CSC gene trees, and the elimination process terminated with a neighbourhood of size 1, a species root partition separating the Epsilonproteobacteria species from the other proteobacteria classes. This LCA is indeed the most frequent one among the CSC gene trees, but with a low frequency of only one in three gene trees. It is noteworthy that the order of elimination does not generally follow the frequency of partitions in the CSC set. For example, the last alternative to be rejected (number 19) was inferred as a root branch in just one tree where it is tied with two other branches, whereas the second and third most frequent CSC roots are rejected already at iterations 19 and 20.

**Figure 5.**
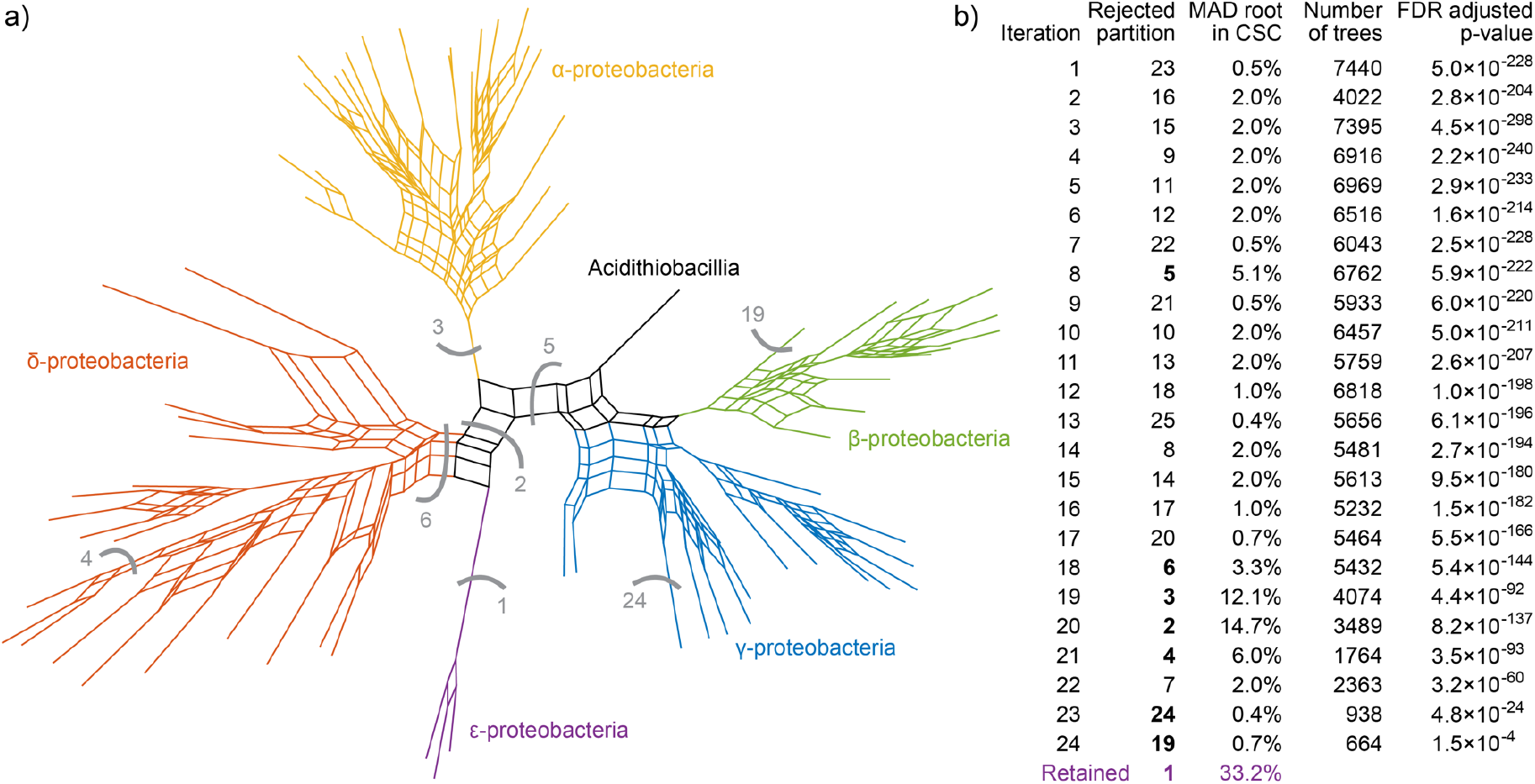
LCA inference by sequential elimination in the proteobacteria dataset. **a)** Phylogenetic split network of the 50 CSC gene trees; **b)** Trace of the sequential elimination process (See Supplementary Table 1b for candidate partition definitions). Selected partitions are indicated by grey arcs in **a)** and bold numbers in **b)**.

The elimination order is determined by the p-value of the one-to-many test, which in turn reflects both the effect size of worse support and the power of the test, where the latter is a function of sample size. Hence, candidate partitions for which a smaller number of gene trees are informative are more difficult to reject. In particular, the testing of an LCA hypothesis of a single basal species partitioned from the other species is limited to those gene families that include the basal species. The last two partitions rejected in Figure 5 are indeed single species partitions, and the number of gene trees that are informative relative both to them and to the remaining candidates drops drastically in comparison to earlier iterations. Yet, even at the last iteration the number of gene trees that bear upon the conclusion is an order of magnitude larger than the number of CSC gene families.

The full complement of the proteobacteria dataset consists of 9,686 gene families. The final conclusion - determination of a single LCA partition - is arrived at by extracting ancestor-descendant information from 86% of the gene families. The gene families that do not provide any evidence consist of 1113 PSC gene families, mainly very small ones (e.g., due to recent gene origin), and 214 PMC families, mostly small families and some with abundant paralogs (e.g., due to gene duplication prior to the LCA).

In the current analysis we used the ancestor deviation measure to provide the strength of the root signal in gene tree branches. We note that the statistical frameworks can accommodate other measures that quantify the root signal for all gene tree branches. A fundamental element in our approach is the prior definition of a pool of candidate root partitions. We advocate deriving the initial set from roots inferred for CSC gene trees. A yet larger but manageable initial set may be constructed of splits frequently observed in the CSC gene trees. Importantly, the initial set need not be limited to observed partitions, but can be augmented by *a-priori* hypotheses informed by current phylogenetic and taxonomical precepts.

The inferred species root partition for the proteobacteria dataset indicates that the proteobacteria LCA was more closely related to modern Epsilonproteobacteria in comparison to the other classes; characteristics of present-day species in that group can therefore be used to hypothesize about the biology of the proteobacterial LCA. Epsilonproteobacteria species show versatile biochemical strategies to fix carbon, enabling members of this class to colonize extreme environments such as deep sea hydrothermal vents (Campbell et al. 2006). Epsilonproteobacteria residing in deep-sea habitats are generally anaerobes and many are characterized as chemolithoautotrophs (Takai et al. 2005). The possibility that the ancient proteobacteria LCA had a chemolithoautotrophic and anaerobic life-style for ancient lineages is in line with the scenario of life’s early phase as predicted by the hydrothermal-vent theory for the origin of life (Martin et al. 2008).

## Discussion

From a purely theoretical perspective, the inference of the LCA for a group of species amounts to the reconstruction of just one branch - the root branch - of the true unrooted species tree, and should therefore be a much easier task than the full resolution of the rooted species tree. Approaches that pose the LCA problem in terms of rooting of a resolved species tree require the solution of a much harder problem as a prerequisite for addressing the easier task. Methods where the input information passes through a bottleneck of a single inferred species tree have the disadvantage that the actual inference of the LCA is based on a sample of size one.

An alternative approach for rooting species trees has emerged from the use of ‘gene-tree-species-tree reconciliation’ models (or simply tree reconciliations) (Szöllősi et al. 2012; Williams et al. 2017; Coleman et al. 2021). These models operate on topological differences between gene-trees and a species tree and the analyses attempt to bring the tree differences into agreement by invoking gene transfers, gene duplications, and gene losses, in any combination as needed. To discern among alternative reconciliation scenarios the evolutionary rates for gene transfers, gene duplications and gene losses need to be estimated from the data. Rate estimation is, however, challenging and a previous study showed that evolutionary rates automatically estimated by a popular tree reconciliation model are often unrealistic and that the use of incorrect rates leads to incorrect root inferences (Bremer et al. 2022). Indeed, evolutionary rates as estimated by tree reconciliation analyses often contradict more conservative estimates obtained by independent studies (Treangen and Rocha 2011; Tria and Martin 2021), which may indicate possible biases within reconciliation models. By contrast, the rooting approach presented here does not rely on *a priori* estimates of rates of gene duplication and transfer and, as such, it offers a simpler solution to the species root problem. Furthermore, and contrary to tree reconciliations, our approach can perform rooting when the species tree is altogether unknown (albeit a candidate root partition is still necessary), which makes our tests especially appealing to rooting prokaryotic phylogenies for which species tree inferences are typically challenging.

Avoiding the reliance on a species tree prompt us to re-evaluate what phylogenetic signal is directly relevant to LCA inference, and to recast the task as that of sampling the total evidence from all gene families at the genomic scope. Moreover, dispensing with a single rooting operation of a single species tree facilitates the reformulation of LCA and root inference in the framework of statistical hypothesis testing. The analytical procedure we outline allows formally to test competing *a-priori* LCA hypotheses and to infer confidence sets for the earliest speciation events in the history of a group of species. Indeed, our approach relies on the ancestor deviation measure that may be sensitive to large deviations from clock-like evolutionary rates. Nonetheless, previous studies suggested that the majority of protein families in prokaryotic genomes are characterized by clock-like evolutionary rates (Novichkov et al. 2004; Dagan et al. 2010). Additionally, we propose that the use of large gene tree samples assists in overcoming bias in the root inference that are due to conflicting signals in the data. Possible biases in the inference of species root partition may arise due to methodological artifacts (for example, tree reconstruction errors) and confounding evolutionary processes such as lateral gene transfer, gene duplication and gene loss. Indeed, our root inferences were consistent across samples of different gene tree categories (CSC, CMC, PSC and PMC), each category bearing different types and degrees of conflicting signals. For instance, gene duplications are more frequent in multicopy gene trees (CMC and PMC), whereas gene losses are likely more frequent in partial trees (PSC and PMC). Phylogenomic inferences utilizing all gene trees for a species set is expected to increase the robustness of the root inference.

Our analyses of the demonstrative datasets show that different species sets present varying levels of LCA signal: the opisthokonta dataset shows a strong root signal, the proteobacteria dataset has a moderate LCA signal. Datasets with weak signal are better described in terms of a confidence sets for root partitions, reflecting the inherent uncertainties and avoiding the pitfalls in forcing a single-hypothesis result.

The LCA inferences presented here utilized 43-86% of the total number of gene families for root partition inferences. This is in stark contrast to the 0.5-1.3% of the gene families that are CSC and can be utilized by traditional approaches. The number of genes families considered in our tests corresponds to the number of genes encoded in modern genomes, supplying ‘total evidence’ for LCA inferences.

## Data & Methods

Protein families for the opisthokonta and proteobacteria datasets were extracted from EggNOG version 4.5 (Huerta-Cepas et al. 2016). Protein families were filtered based on the number of species, gene copy number, number of OTUs, and sequence length, as follows. Protein families present in less than four species were discarded. Suspected outlier sequences were detected based on their length relative to the median length: sequences were removed if shorter than half or longer than twice the median. Species with more than ten copies of a gene were removed from the corresponding gene family. Multi-copy gene families were discarded if the number of species was smaller than half the total number of OTUs (Table 1).

Protein sequences of the resulting protein families were aligned using MAFFT version 7.027b with L-INS-i alignment strategy (Katoh and Standley 2013). Phylogenetic trees were reconstructed using iqtree version 1.6.6 with the model selection parameters ‘-mset LG-madd LG4X’ (Nguyen et al. 2015). The phylogenetic network (Fig. 5) was reconstructed using SplitsTree4 version 4.14.6 (Huson and Bryant 2006). Branch ancestral deviation (AD) values and roots for the consensus analysis were inferred using mad.py version 2.21 (Tria et al. 2017).

The source code to run the phylogenomic rooting as well as the unrooted trees with AD values used in this study are found in the following repository: https://github.com/deropi/PhyloRooting.git. Additionally, R code to replicate some of the figures in this paper is also provided.

## Supplementary Information

Supplementary tables 1 and 2 are available online at: https://doi.org/10.6084/m9.figshare.17156030

## Acknowledgements

We thank Maxime Godfroid for fruitful discussions. The study was supported by CAPES (Coordination for the Improvement of Higher Education Personnel–Brazil) (awarded to FDKT) and the European Research Council (Grant No. 281357 awarded to TD). DRP is supported by the DFG funded CRC1182 the origin and function of metaorganisms.

## Notes

https://doi.org/10.6084/m9.figshare.17156030

### Competing Interest Statement

The authors have declared no competing interest.

### Summary of Updates

Minor referee corrections

## References

Benjamini Y., Hochberg Y. 1995. Controlling the false discovery rate: A practical and powerful approach to multiple testing. J R Stat Soc Series B Stat Methodol. 57:289–300.

Bettisworth B., Stamatakis A. 2021. Root Digger: a root placement program for phylogenetic trees. BMC Bioinformatics. 22:225.

Bremer N., Knopp M., Martin W.F., Tria F.D.K. 2022. Realistic Gene Transfer to Gene Duplication Ratios Identify Different Roots in the Bacterial Phylogeny Using a Tree Reconciliation Method. Life. 12:995.

Campbell B.J., Engel A.S., Porter M.L., Takai K. 2006. The versatile ε-proteobacteria: key players in sulphidic habitats. Nat Rev Microbiol. 4:458–468.

Cherlin S., Heaps S.E., Nye T.M.W., Boys R.J., Williams T.A., Embley T.M. 2018. The Effect of Nonreversibility on Inferring Rooted Phylogenies. Molecular Biology and Evolution. 35:984–1002.

Ciccarelli F.D., Doerks T., von Mering C., Creevey C.J., Snel B., Bork P. 2006. Toward automatic reconstruction of a highly resolved tree of life. Science. 311:1283–1287.

Coleman G.A., Davín A.A., Mahendrarajah T.A., Szánthó L.L., Spang A., Hugenholtz P., Szöllősi G.J., Williams T.A. 2021. A rooted phylogeny resolves early bacterial evolution. Science. 372:eabe0511.

Dagan T., Martin W. 2006. The tree of one percent. Genome Biol. 7:118.

Dagan T., Roettger M., Bryant D., Martin W. 2010. Genome networks root the tree of life between prokaryotic domains. Genome Biology and Evolution. 2:379–392.

Dagan T., Roettger M., Stucken K., Landan G., Koch R., Major P., Gould S.B., Goremykin V. V., Rippka R., Tandeau de Marsac N., Gugger M., Lockhart P.J., Allen J.F., Brune I., Maus I., Pühler A., Martin W.F. 2013. Genomes of Stigonematalean cyanobacteria (Subsection V) and the evolution of oxygenic photosynthesis from prokaryotes to plastids. Genome Biol Evol. 5:31–44.

Doolittle W.F., Bapteste E. 2007. Pattern pluralism and the Tree of Life hypothesis. Proc Natl Acad Sci USA. 104:2043–2049.

Eisen J.A. 2003. Phylogenomics: Intersection of Evolution and Genomics. Science. 300:1706–1707.

Farris J.S. 1972. Estimating Phylogenetic Trees from Distance Matrices. Am Nat. 106:645–668.

Fitch W.M., Margoliash E. 1967. Construction of phylogenetic trees. Science. 155:279–284.

Fox G.E., Stackebrandt E., Hespell R.B., Gibson J., Maniloff J., Dyer T.A., Wolfe R.S., Balch W. E., Tanner R.S., Magrum L.J., Zablen L.B., Blakemore R., Gupta R., Bonen L., Lewis B.J., Stahl D.A., Luehrsen K.R., Chen K.N., Woese C.R. 1980. The phylogeny of Prokaryotes. Science. 209:457–463.

Gogarten J.P., Kibak H., Dittrich P., Taiz L., Bowman E.J., Bowman B.J., Manolson M.F., Poole R.J., Date T., Oshima T., Konishi J., Denda K., Yoshida M. 1989. Evolution of the vacuolar H+-ATPase: implications for the origin of eukaryotes. Proceedings of the National Academy of Sciences. 86:6661–6665.

Hammerschmidt K., Landan G., Domingues Kümmel Tria F., Alcorta J., Dagan T. 2021. The order of trait emergence in the evolution of cyanobacterial multicellularity. Genome Biol Evol. 13.

Huelsenbeck J.P., Bollback J.P., Levine A.M. 2002. Inferring the Root of a Phylogenetic Tree. Systematic Biology. 51:32–43.

Huerta-Cepas J., Szklarczyk D., Forslund K., Cook H., Heller D., Walter M.C., Rattei T., Mende D.R., Sunagawa S., Kuhn M., Jensen L.J., von Mering C., Bork P. 2016. eggNOG 4.5: a hierarchical orthology framework with improved functional annotations for eukaryotic, prokaryotic and viral sequences. Nucleic Acids Res. 44:D286–D293.

Hug L.A., Baker B.J., Anantharaman K., Brown C.T., Probst A.J., Castelle C.J., Butterfield C.N., Hernsdorf A.W., Amano Y., Ise K., Suzuki Y., Dudek N., Relman D.A., Finstad K.M., Amundson R., Thomas B.C., Banfield J.F. 2016. A new view of the tree of life. Nat Microbiol. 1:1–6.

Huson D.H., Bryant D. 2006. Application of phylogenetic networks in evolutionary studies. Mol Biol Evol. 23:254–267.

Iwabe N., Kuma K., Hasegawa M., Osawa S., Miyata T. 1989. Evolutionary relationship of archaebacteria, eubacteria, and eukaryotes inferred from phylogenetic trees of duplicated genes. Proc Natl Acad Sci U S A. 86:9355–9359.

Katoh K., Standley D.M. 2013. MAFFT multiple sequence alignment software version 7: Improvements in performance and usability. Mol Biol Evol. 30:772–780.

Katz L.A. 2012. Origin and Diversification of Eukaryotes. Annu Rev Microbiol. 66:411–427.

Kluge A.G., Farris J.S. 1969. Quantitative Phyletics and the Evolution of Anurans. Syst Biol. 18:1–32.

Lang J.M., Darling A.E., Eisen J.A. 2013. Phylogeny of bacterial and archaeal genomes using conserved genes: supertrees and supermatrices. PLoS One. 8:e62510–15.

Lepage T., Bryant D., Philippe H., Lartillot N. 2007. A General Comparison of Relaxed Molecular Clock Models. Mol Biol Evol. 24:2669–2680.

Linz S., Radtke A., von Haeseler A. 2007. A likelihood framework to measure horizontal gene transfer. Mol Biol Evol. 24:1312–1319.

Lovejoy C.O., Suwa G., Simpson S.W., Matternes J.H., White T.D. 2009. The great divides: *Ardipithecus ramidus* reveals the postcrania of our last common ancestors with african apes. Science. 326:100–106.

Mai U., Sayyari E., Mirarab S. 2017. Minimum variance rooting of phylogenetic trees and implications for species tree reconstruction. PLOS ONE. 12:e0182238.

Martin W., Baross J., Kelley D., Russell M.J. 2008. Hydrothermal vents and the origin of life. Nat Rev Microbiol. 6:805–814.

Medini D., Donati C., Tettelin H., Masignani V., Rappuoli R. 2005. The microbial pan-genome. Curr Opin Genet Dev. 15:589–594.

Morel B., Schade P., Lutteropp S., Williams T.A., Szöllősi G.J., Stamatakis A. 2022. SpeciesRax: A Tool for Maximum Likelihood Species Tree Inference from Gene Family Trees under Duplication, Transfer, and Loss. Molecular Biology and Evolution. 39:msab365.

Naser-Khdour S., Quang Minh B., Lanfear R. 2022. Assessing Confidence in Root Placement on Phylogenies: An Empirical Study Using Nonreversible Models for Mammals. Systematic Biology. 71:959–972.

Nguyen L.-T., Schmidt H.A., von Haeseler A., Minh B.Q. 2015. IQ-TREE: A fast and effective stochastic algorithm for estimating maximum-likelihood phylogenies. Mol Biol Evol. 32:268–274.

Novichkov P.S., Omelchenko M.V., Gelfand M.S., Mironov A.A., Wolf Y.I., Koonin E.V. 2004. Genome-Wide Molecular Clock and Horizontal Gene Transfer in Bacterial Evolution. Journal of Bacteriology. 186:6575–6585.

Okamoto E., Kusakabe R., Kuraku S., Hyodo S., Robert-Moreno A., Onimaru K., Sharpe J., Kuratani S., Tanaka M. 2017. Migratory appendicular muscles precursor cells in the common ancestor to all vertebrates. Nat Ecol Evol. 1:1731–1736.

Parks D.H., Chuvochina M., Waite D.W., Rinke C., Skarshewski A., Chaumeil P.-A., Hugenholtz P. 2018. A standardized bacterial taxonomy based on genome phylogeny substantially revises the tree of life. Nat Biotechnol. 36:996–1004.

Pisani D., Cotton J.A., McInerney J.O. 2007. Supertrees disentangle the chimerical origin of eukaryotic genomes. Mol Biol Evol. 24:1752–1760.

Semple C., Steel M. 2001. Tree reconstruction via a closure operation on partial splits. Computational Biology.:126–134.

Smith M.L., Hahn M.W. 2021. New Approaches for Inferring Phylogenies in the Presence of Paralogs. Trends in Genetics. 37:174–187.

Stechmann A., Cavalier-Smith T. 2002. Rooting the eukaryote tree by using a derived gene fusion. Science. 297:89–91.

Swenson K.M., El-Mabrouk N. 2012. Gene trees and species trees: irreconcilable differences. BMC Bioinformatics. 13:S15.

Szöllősi G.J., Boussau B., Abby S.S., Tannier E., Daubin V. 2012. Phylogenetic modeling of lateral gene transfer reconstructs the pattern and relative timing of speciations. Proc Natl Acad Sci USA. 109:17513–17518.

Szöllősi G.J., Tannier E., Daubin V., Boussau B. 2015. The inference of gene trees with species trees. Syst Biol. 64:e42–e62.

Takai K., Campbell B.J., Cary S.C., Suzuki M., Oida H., Nunoura T., Hirayama H., Nakagawa S., Suzuki Y., Inagaki F., Horikoshi K. 2005. Enzymatic and genetic characterization of carbon and energy metabolisms by deep-sea hydrothermal chemolithoautotrophic isolates of *Epsilonproteobacteria*. Appl Environ Microbiol. 71:7310–7320.

Treangen T.J., Rocha E.P.C. 2011. Horizontal Transfer, Not Duplication, Drives the Expansion of Protein Families in Prokaryotes. PLoS Genet. 7:e1001284–12.

Tria F.D.K., Landan G., Dagan T. 2017. Phylogenetic rooting using minimal ancestor deviation. Nat Ecol Evol. 1:0193.

Tria F.D.K., Martin W.F. 2021. Gene Duplications Are At Least 50 Times Less Frequent than Gene Transfers in Prokaryotic Genomes. Genome Biol Evol. 13:evab224.

Waite D.W., Vanwonterghem I., Rinke C., Parks D.H., Zhang Y., Takai K., Sievert S.M., Simon J., Campbell B.J., Hanson T.E., Woyke T., Klotz M.G., Hugenholtz P. 2017. Comparative genomic analysis of the Class *Epsilonproteobacteria* and proposed reclassification to Epsilonbacteraeota (phyl. nov.). Front Microbiol. 8:4962–19.

Weiss M.C., Sousa F.L., Mrnjavac N., Neukirchen S., Roettger M., Nelson-Sathi S., Martin W.F. 2016. The physiology and habitat of the last universal common ancestor. Nat Microbiol.:1–8.

Whidden C., Zeh N., Beiko R.G. 2014. Supertrees Based on the Subtree Prune-and-Regraft Distance. Syst Biol. 63:566–581.

Williams T.A., Cox C.J., Foster P.G., Szöllősi G.J., Embley T.M. 2020. Phylogenomics provides robust support for a two-domains tree of life. Nat Ecol Evol. 4:138–147.

Williams T.A., Heaps S.E., Cherlin S., Nye T.M.W., Boys R.J., Embley T.M. 2015. New substitution models for rooting phylogenetic trees. Philos Trans R Soc B: Biol Sci. 370:20140336–8.

Williams T.A., Szöllősi G.J., Spang A., Foster P.G., Heaps S.E., Boussau B., Ettema T.J.G., Embley T.M. 2017. Integrative modeling of gene and genome evolution roots the archaeal tree of life. Proc Natl Acad Sci USA. 114:E4602–E4611.

